# Human Connectome Project resting state fMRI data organized into 60 fine-scaled retinotopically organized regions in cortical areas V1, V2 and V3

**DOI:** 10.1101/340174

**Authors:** Debra Ann Dawson, Zixuan Yin, Jack Lam, Amir Shmuel

## Abstract

The data comprises 60 regions of interest (ROIs) from V1, V2, and V3 of the human visual cortex. Preprocessed data from the Human Connectome Project (HCP) 900 subjects public data release were utilized: 220 subjects were randomly selected, each with 4 scans of resting state fMRI data. Given that these subjects did not have retinotopy scans performed, the visual areas were defined using an anatomical template from Benson et al. (2014). Visual areas from each hemisphere were further divided along dorsal-ventral lines into quadrants, resulting in 4 quadrants per subject. Within each quadrant, fine scaled ROIs were defined by subdividing each visual area into 5 regions according to eccentricity. These data may be useful for studying retinotopically organized functional connectivity in the visual cortex using the HCP 3 Tesla dataset.

## Specifications Table

|  |  |
| --- | --- |
| Subject area | *Neuroscience* |
| More specific subject area | *NeuroImaging* |
| Type of data | *Functional MRI (fMRI) region of interest average time courses.* |
| How data was acquired | *fMRI at 3 Tesla, Human Connectome Project public release data.* |
| Data format | *Voxel time courses averaged into regions of interest; Matlab array.* |
| Experimental factors | *Template of early visual areas applied to preprocessed data* |
| Experimental features |  |
| Data source location | *Human Connectome Project public data release* |
| Data accessibility | *Data is with this article (with Data In Brief)* |
| Related research article | Dawson DA, Lam J, Lewis LB, Carbonell F, Mendola JD, Shmuel A. Partial Correlation-Based Retinotopically Organized Resting-State Functional Connectivity Within and Between Areas of the Visual Cortex Reflects More Than Cortical Distance. *Brain Connect.* Feb 2016;6(1):57-75. |

## Value of the Data

- Most datasets of subjects having undergone retinotopy include data from only a small number of subjects, limiting the power possible from the study. Here we present high quality data obtained from a large number (220) of subjects.
- The data can be used for studying the functional connectivity within and between areas V1, V2 and V3 of the visual cortex.
- Most of the previously acquired data from study subjects having undergone retinotopy scanning has been done with a 3 Tesla scanner. We provide a large, high quality sample of V1, V2, and V3 resting state data acquired from a 3 Tesla scanner, allowing for the opportunity to undertake comparative studies using other datasets, including datasets from patient populations (e.g. Mendola et al., 2018).

## Data

The data comprises 60 regions of interest (ROIs) from V1, V2, and V3 of the human visual cortex. 3D volume preprocessed resting state fMRI data from the Human Connectome Project (HCP) 900 subjects public data release were utilized: 220 random subjects were selected, each with 4 scans of 3 Tesla resting state fMRI data. Given that these subjects did not have retinotopy scans performed, the visual areas were defined using an anatomical template from Benson et al. (2014). Visual areas from each hemisphere were further subdivided along dorsal-ventral lines into quadrants, resulting in 4 quadrants per subject. Within each quadrant, fine scaled ROIs were defined by subdividing each visual area into 5 regions according to eccentricity.

## Experimental Design, Materials, and Methods

### Original data and preprocessing

Subjects were randomly selected from those available in the HCP900 data release. 3D volume preprocessed resting state fMRI data were used, along with files resulting from a FreeSurfer-based surface extraction pipeline. Important details regarding scanning follows (see also Table 1): the resting state fMRI scans were done with 3 Tesla scanners, the TR was 0.72s, there are 1200 time points per scan. Full details on the data acquisition can be found in a special Connectome issue of NeuroImage, the most important contributions to the data used here being the following: Van Essen et al., 2013, Ŭgurbil et al., 2013, Glasser et al. 2013, Smith et al., 2013, Sotiropoulos et al., 2013.

**Table 1.** Parameters relevant for the resting state fMRI images from the HCP dataset. This table is taken directly from the HCP 900 Subjects Data Release Reference Manual (page 32). https://www.humanconnectome.org/storage/app/media/documentation/s900/HCP_S900_Rel_ease_Reference_Manual.pdf

| Parameter | Value |
| --- | --- |
| Sequence | Gradient-echo EPI |
| TR | 720 ms |
| TE | 33.1 ms |
| flip angle | 52 deg |
| FOV | 208x180 mm (RO x PE) |
| Matrix | 104x90 (RO x PE) |
| Slice thickness | 2.0 mm; 72 slices; 2.0 mm isotropic voxels |
| Multiband factor | 8 |
| Echo spacing | 0.58 ms |
| BW | 2290 Hz/Px |

The preprocessed HCP data have undergone “the Minimal Preprocessing Pipelines”. The specific pipelines the data utilized here underwent are the so called PreFreesurfer, Freesurfer, PostFreeSurfer, and fMRIVolume pipelines. Details on these and all of the Minimal Preprocessing Pipelines can be found in Glasser et al. (2013), with additional details in Jenkinson et al. (2012), Fischl (2012), and Jenskinson et al. (2002). These pipelines utilize tools From FSL and FreeSurfer.

The preprocessed data downloaded for this data release came from the following HCP repository locations:

~~~
/HCP_900/{subjectID}/T1w/{subjectID}/*
/HCP_900/{subjectID}/MNINonLinear/Results/rfMRI_REST{sessionID}_{runID}/…
rfMRI_REST{sessionID}_{runID}_hp2000_clean.nii.gz
~~~

The first contains the post-FreeSurfer brain extracted volumes and surfaces which are needed for ROI definition. The second contains the four resting state scans per subject, with {sessionID} holding the value of either 1, or 2, and {runID} holding the value of either LR or RL (this is meant to represent the phase encoding direction of the oblique axial acquisitions, alternated in consecutive runs: LR=left-right, RL=right-left). {subjectID} is the six digit subject number assigned to all participants in the HCP.

**Figure 1.**
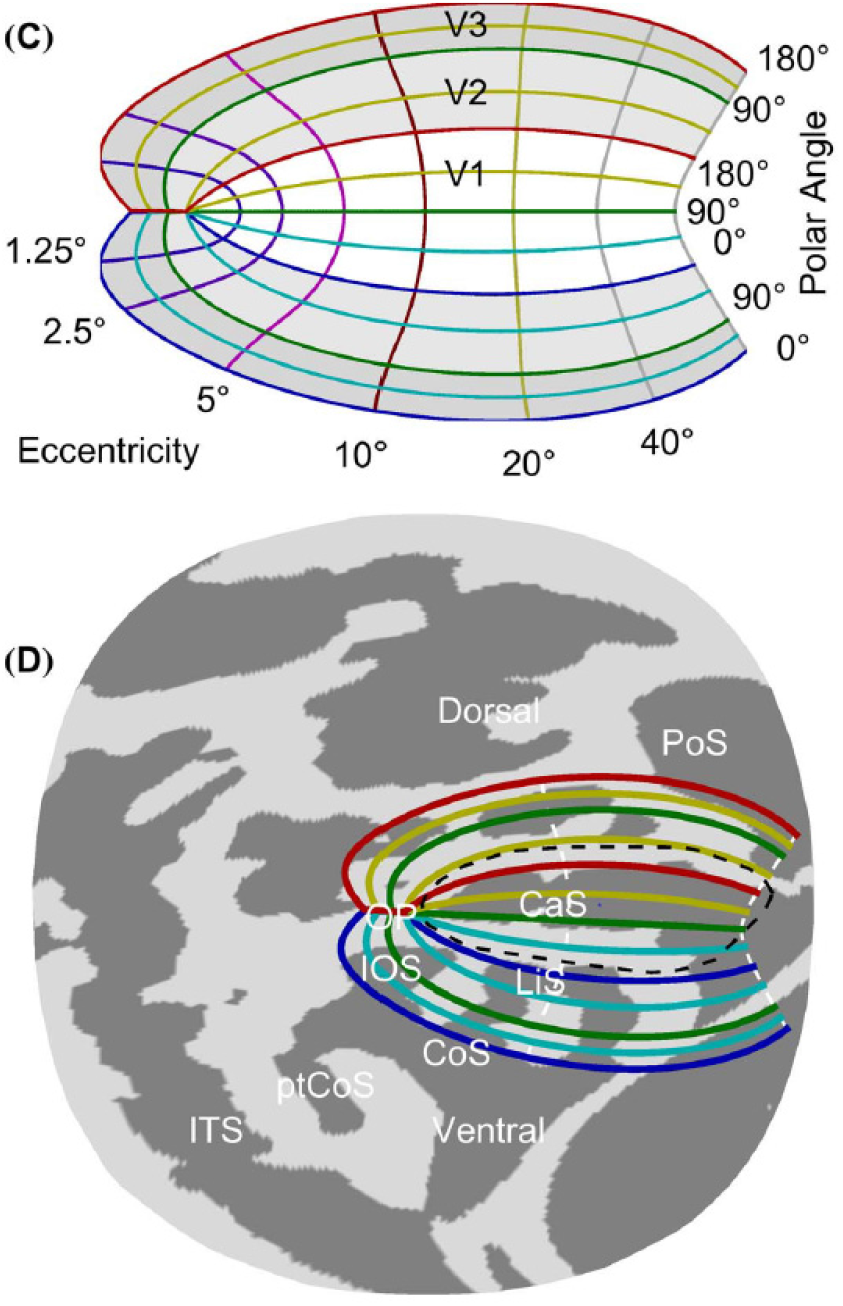
Figure (panels C and D) from Benson et al. (2014). “(C) The algebraic model of retinotopic organization. V1, V2, and V3 are colored white, light gray, and dark gray, respectively. (D) The cortical surface atlas space (fsaverage_sym) from the occipital pole after flattening to the 2D surface. The Hinds V1 border (Hinds et al., 2008) is indicated by the dashed black line. The algebraic model of retinotopic organization used in registration is plotted with all 0°, 90°, and 180° polar angle lines colored according to their color in (C), and the 10° and 90° eccentricity lines dashed and colored white. Shown are the Calcarine Sulcus (CaS), the Parietal-occipital Sulcus (PoS), the Lingual sulcus (LiS), the Inferior Occipital Sulcus (IOS), the Collateral Sulcus (CoS), the posterior Collateral Sulcus (ptCoS), the Inferior Temporal Sulcus (ITS), and the Occipital Pole (OP).” Reprinted from Benson et al. (2014) with permission.

**Figure 2.**
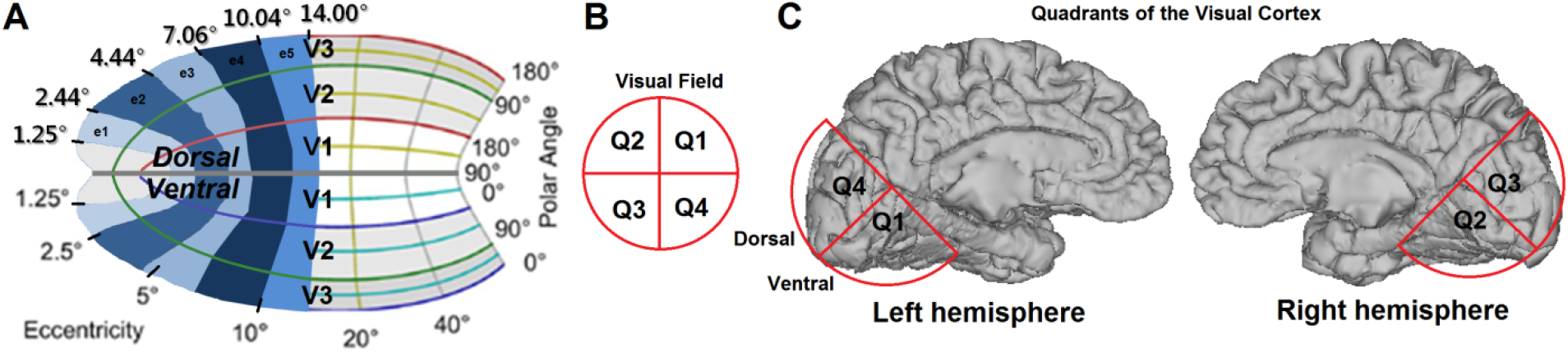
ROIs defined on a template representation of one hemisphere of the visual cortex. (A) We define the regions using the template from Benson et al. (2014). We first split each hemisphere along dorsal-ventral lines, forming two quadrants per hemisphere. Voxels from each visual area and quadrant were classified into five eccentricity bins with the boundaries 1.25° to 2.44°, 2.44° to 4.44°, 4.44° to 7.06°, 7.06° to 10.04°, and 10.04° to 14.00°. Panel (C) presents an approximate scheme of the four quadrants. These quadrants reflect the visual field representations as identified in (B). For preparing Figure 2A, we modified Figure 1C from Benson et al (2014) with permission.

**Figure 3.**
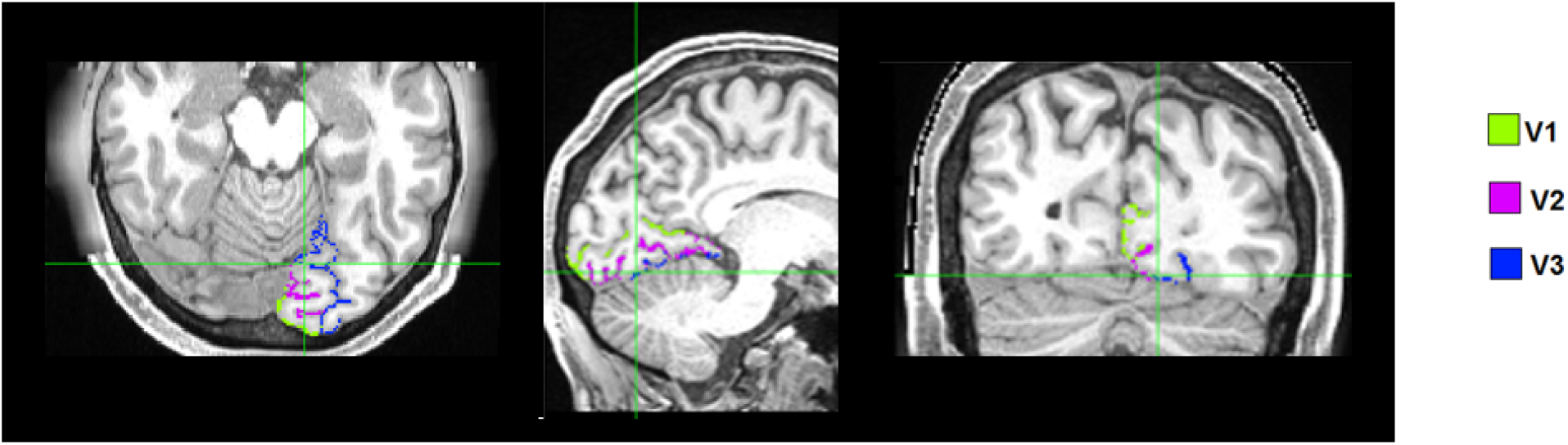
Results of template alignment on an example subject, ID 1004008. The green outline indicates the spanned stretch of the cortical surface assigned to V1, purple is V2, and blue is V3.

### Regions of interest

We define a local visual network within a hemisphere, which is subdivided into ventral and dorsal sections along the horizontal meridian of the polar angle phase map (>90° = dorsal quadrant, <90° = ventral quadrant, see figure 2A). Anatomically, this horizontal meridian is expected to lie at the deepest curvature of the calcarine sulcus (figure 2C). These data were obtained from the two hemispheres of each subject, thus forming 4 quadrants of the visual cortex (cortical regions which respond to stimuli in the 4 quadrants of the visual field) (Figure 2C).

Typically, visual areas are defined on the cortex via retinotopic scanning and mapping. Rotating wedge and expanding/contracting ring stimuli are presented to the subjects during a retinotopic scanning session in order to elicit polar angle and eccentricity phase maps respectively. Visual area delineation is achieved by examining the reversal of direction of increasing polar angle phases perpendicular to the iso-eccentricity lines represented on the cortex (Sereno et al., 1995; Wandell et al., 2007).

We were interested in having access to the broad range of subjects who were only scanned in a 3 Tesla scanner and thus sought a method which would allow us to define the visual areas without having retinotopic scans. As a result, we have delineated V1, V2, and V3 using a template (version 2.5) described by Benson et al. (2012, 2014). The template was produced by aligning aggregate retinotopy from a group of subjects to a model of V1, V2, and V3 retinotopy based on surface topology in the visual cortex (Figure 2). We applied this template to the HCP subjects’ *orig.mgz* (output file from Freesurfer’s surface extraction) after computing the registration of the template to the subject (Figure 3). The registration is accomplished via alignment of the *sphere.reg* subject file to *fsaverage_sym.sphere.reg,* an average surface atlas available in Freesurfer to which the template is already aligned.

ROIs are subsections (5) of visual areas V1, V2, and V3 (3) within each quadrant (4) of the visual cortex (total number of ROIs=5×3×4 = 60) (Figure 2). Quadrants are defined by separating the visual cortex into hemispheres, and then along dorsal/ventral lines according to polar phase. These quadrants as defined on the cortex respond to a quadrant of the visual field. It can be very problematic in FC analyses if voxels belonging to one functional region are misclassified into a neighboring region (Smith et al., 2011). In order to avoid this detrimental effect, we removed a 10 polar angle degree range from either side of the border between visual areas. We also excluded eccentricity values below 1.25° due to measurement bias near the edge of the stimulus range and difficulty in obtaining eccentricity mapping near the fovea (Wandell et al., 2007; Schira et al., 2009; Benson et al., 2014). The retinotopy template assigns values to voxels using the full extent of possible phases in eccentricity and polar angle dimensions, i.e. 0° to 90°. This range of eccentricity values goes beyond what is typical in experimental retinotopic mapping (where the maximum eccentricity of a stimulus presented is usually under 15°). We therefore have selected a more “realistic” maximal eccentricity boundary for use in defining our ROIs. We chose our maximal eccentricity to be 14°. This value is the same as we had used in a previous study by our group with measured retinotopy (Dawson et al., 2016) and is within the range of values tested in the Benson et al. (2014) study which defined the template.

In Benson et al., they look predominantly at an eccentricity range of 0° to10° (avoiding 1.25° at each edge), but also considered a maximum of 20°. They found a somewhat decreased match between template predicted visual area boundaries and measured retinotopic boundaries with a maximum eccentricity of 20° relative to 10°, however the results were still relevant and the errors similar in scale. Importantly, the authors point out that the error in visual area boundary definition between two retinotopic scans is greater than the prediction error of their anatomical template (as compared to the measured retinotopy). Thus, for maximal eccentricity values under 20°, the discordance of the template with measured retinotopic visual area delineation is expected to be within the range of the uncertainty of the measured retinotopy. Our choice of 14° is a compromise in order to capture a broader field while minimizing prediction error.

Given the above choices regarding upper and lower limits of eccentricity, we selected boundaries along the eccentricity dimension within each visual area, delineating our fine-scaled ROIs as follows: 1.25° to 2.44°, 2.44° to 4.44°, 4.44° to 7.06°, 7.06° to 10.04°, and 10.04° to 14.00°. The middle boundaries were chosen via an optimization algorithm in order to approximately equalize the number of voxels included in each ROI across subjects. The eccentricity ranges of the regions are expected to increase as we move more peripherally due to the smaller cortical magnification in peripheral representations of the visual field relative to foveal representations (Duncan and Boynton, 2003). We denote these five regions as e1, e2, e3, e4, and e5, respectively (Figure 2A). ‘e1’ is an abbreviation for ‘eccentricity 1’, representing the cortical region which responds to stimuli most central in the visual field. ‘e5’ represents the region which responds to stimuli most peripheral in the visual field. ROI time courses are generated by computing the mean across all voxels within the ROI. These average time courses have been normalized by subtracting the mean and dividing by the standard deviation across time points.

### Details of data organization

DATAHCP200 is a 1×880 cell Matlab array. Each cell represents an independent run. The order of the runs is as follows:

~~~
Subject1_run1_LR, Subject1_run1_RL, Subject1_run2_LR, Subject1_run2_RL,
Subject2_run1_LR, Subject2_run1_RL, Subject2_run2_LR,…, Subject220_run2_RL
~~~

Within each cell is a 1200×60 double matrix. Column vectors contain the time-courses (1200 time points) for each of the 60 ROIs. The TR of these resting state data sets is 0.72s. The order of the ROIs is as follows:

~~~
Q1_V1_e1, Q1_V1_e2, Q1_V1_e3, Q1_V1_e4, Q1_V1_e5, Q1_V2_e1, Q1_V2_e2, …, Q1_V3_e5, Q2_V1_e1,…, Q4_V3_e5.
~~~

Where “Qi” represents the quadrant (i=1,2,3,4), “Vj” represents the visual area (j=1,2,3), and “ek” represents the eccentricity region (k=1, 2, 3, 4, 5) (e1 is most central, e5 is most peripheral; see figure 2A). Quadrant numbering is as presented in Figure 2B and C.

## Acknowledgments

Data were provided by the Human Connectome Project, WU-Minn Consortium (Principal Investigators: David Van Essen and Kamil Ugurbil; 1U54MH091657) funded by the 16 NIH Institutes and Centers that support the NIH Blueprint for Neuroscience Research; and by the McDonnell Center for Systems Neuroscience at Washington University.

## References

1. Benson NC, Butt OH, Brainard DH, Aguirre GK. Correction of distortion in flattened representations of the cortical surface allows prediction of V1-V3 functional organization from anatomy. PLoS Comput Biol. Mar 2014;10(3):e1003538.

2. Benson NC, Butt OH, Datta R, Radoeva PD, Brainard DH, Aguirre GK. The retinotopic organization of striate cortex is well predicted by surface topology. Curr Biol. Nov 6 2012;22(21):2081–2085.

3. Dawson DA, Cha K, Lewis LB, Mendola JD, Shmuel A. Evaluation and calibration of functional network modeling methods based on known anatomical connections. NeuroImage. Feb 15 2013;67:331–343.

4. Dawson DA, Lam J, Lewis LB, Carbonell F, Mendola JD, Shmuel A. Partial Correlation-Based Retinotopically Organized Resting-State Functional Connectivity Within and Between Areas of the Visual Cortex Reflects More Than Cortical Distance. Brain Connect. Feb 2016;6(1):57–75.

5. Fischl B. FreeSurfer. NeuroImage. Aug 15 2012;62(2):774–781.

6. Glasser MF, Sotiropoulos SN, Wilson JA, et al. The minimal preprocessing pipelines for the Human Connectome Project. NeuroImage. 2013/10/15/ 2013;80:105–124.

7. Hinds OP, Rajendran N, Polimeni JR, et al. Accurate prediction of V1 location from cortical folds in a surface coordinate system. NeuroImage. Feb 15 2008;39(4):1585–1599.

8. Jenkinson M, Bannister P, Brady M, Smith S. Improved optimization for the robust and accurate linear registration and motion correction of brain images. NeuroImage. Oct 2002;17(2):825–841.

9. Jenkinson M, Beckmann CF, Behrens TE, Woolrich MW, Smith SM. FSL. NeuroImage. Aug 15 2012;62(2):782–790.

10. Mendola JD, Lam J, Rosenstein M, Lewis LB, Shmuel A. Partial Correlation Analysis Reveals Abnormal Retinotopically Organized Functional Connectivity of Visual Areas in Amblyopia. NeuroImage Clinical, 2018. 18:192–201. https://doi.org/10.1016/j.nicl.2018.01.022.

11. Schira MM, Tyler CW, Breakspear M, Spehar B. The Foveal Confluence in Human Visual Cortex. Journal of Neuroscience. 2009; 29: 9050–9058.

12. Sereno M, Dale AM, Reppas JB, Kwong KK, Belliveau JW, Brady TJ, Rosen BR, Tootell RB. Borders of multiple visual areas in humans revealed by functional magnetic resonance imaging. Science. 1995;269(5212):889–893.

13. Smith SM, Karla L. Miller, Gholamreza Salimi-Khorshidi, Matthew Webster, Christian F. Beckmann, Thomas E. Nichols, Joseph D. Ramsey, Mark W. Woolrich. Network modelling methods for FMRI. Neuroimage. 2011;54:875–891.

14. Smith SM, Beckmann CF, Andersson J, et al. Resting-state fMRI in the Human Connectome Project. NeuroImage. 2013/10/15/ 2013;80:144–168.

15. Sotiropoulos SN, Jbabdi S, Xu J, et al. Advances in diffusion MRI acquisition and processing in the Human Connectome Project. NeuroImage. 2013/10/15/ 2013;80:125–143.

16. Uğurbil K, Xu J, Auerbach EJ, et al. Pushing spatial and temporal resolution for functional and diffusion MRI in the Human Connectome Project. NeuroImage. 2013/10/15/ 2013;80:80–104.

17. Van Essen DC, Smith SM, Barch DM, Behrens TEJ, Yacoub E, Ugurbil K. The WU-Minn Human Connectome Project: An overview. NeuroImage. 2013/10/15/ 2013;80:62–79.

18. Wandell BA, Dumoulin SO, Brewer AA. Visual Field Maps in Human Cortex. Neuron. 2007/10/25/ 2007;56(2):366–383.

